# ARCHAEOGENOMIC AND BIOINFORMATIC ANALYSIS OF THE COLUMBUS LINEAGE: EVIDENCE FROM THE COUNTS OF GELVES

**DOI:** 10.64898/2026.04.01.715912

**Authors:** I Navarro-Vera, A Bonilla, M Tirapu, M Albert, P.P Jímenez, D Herranz-Rodrigo, R Cruz Alcázar, M.C García, Yravedra Sainz de los Terreros

## Abstract

The geographical and familial origins of Christopher Columbus have remained a subject of intense historiographical debate for over five centuries. Despite numerous hypotheses, empirical genetic evidence capable of resolving his ancestral history or place of birth has been absent from the literature until now. This study presents the third stage of the first forensic genetic analysis performed on skeletal remains belonging to several direct descendants of Columbus, spanning the 16^th^ to 18^th^ centuries. By applying Massively Parallel Sequencing (MPS) to analyse autosomal, X- and Y- chromosome DNA markers, and integrating the results with multidisciplinary evidence from historical, genealogical, archaeological, and anthropological research implicated in this project, the identification of several individuals founded in the Crypt of Santa María de Gracia located in Gelves (Sevilla, Spain) has been achieved. The analysis of their biological relatedness enabled the reconstruction of kinship networks among the individuals interred in the crypt, which, when interpreted in the context of documented genealogical lineages, provides indirect but consistent evidence pointing toward the debated origin of the discoverer.

## INTRODUCTION

For more than 500 years, the precise origin of Christopher Columbus has been one of history’s most enduring enigmas. The intentional obfuscation of his early life by Columbus himself ^1,2^ has led to a myriad of conflicting theories proposing various European sites as his place of birth. To date, a lack of scientific certainty has persisted due to the absence of conclusive genetic data.

Most studies investigating the origins of Admiral Christopher Columbus are based on circumstantial evidence and indirect findings, which are often influenced by no scientific motivations. However, his origins have rarely been examined from a rigorous scientific perspective. The objective of this study was to analyse the origins of Christopher Columbus from an expert standpoint, employing a multidisciplinary approach and a range of scientific techniques.

To scientifically address the ancestral identity of Christopher Columbus, this study targeted the primary burial site of his direct lineage: the Santa María de Gracia church in Gelves (Seville; 37.33848, -6.02730) (See 3d model of the church) ^3^. The site serves as the pantheon for the Counts of Gelves, housing the largest concentration of Columbus’ direct descendants, at least seven, including his granddaughter. The selection of this crypt as the source for genomic and anthropological samples is justified by several critical factors. The church serves as a monumental testament to the Columbus lineage. The transept dome features a prominent relief of the House of Columbus arms, inscribed with the foundational motto: *“A Castilla y a León nuevo mundo dio Colón”* (*To Castile and León, Columbus gave a New World*). This iconography is further reinforced by the presence of both the Columbus coat of arms and those of the Royal House of Portugal on the four pendentives.

Beyond its symbolic importance, the crypt offers unique analytical advantages due to its unprofaned state and the distinctive cultural markers of its funerary architecture. Situated beneath the high altar, the burial chamber follows a Lusitanian tradition—virtually non-existent in the Spanish context of the period—where coffins are placed in large, vaulted niches (both in hight and width) upon pedestals to prevent contact with the floor, allowing them to be seen through the grilles or doors that lead into the niche. This architectural arrangement underscores the Portuguese identity inherent to this branch of the lineage, aligning with the historical and genealogical records of the family ^4,5,6^. The exceptional preservation of the remains, coupled with precise genealogical traceability, provides a verified biological framework for the high-resolution genomic and isotopic analyses conducted in this research.

Over the past years, an extensive multidisciplinary investigation has been carried out, encompassing genealogical, historical, archaeological, anthropological, and forensic genetic research. Advances in technology in each of these fields have made it possible to obtain diverse datasets, which have helped progress the study and establish a comprehensive line of research with a broad scope.

The skeletal remains of all individuals were commingled and distributed across four unidentified boxes (Box 1–4). Following the anthropological study, the minimum number of individuals (MNI) was established at 12, with biological sex estimation indicating six males and six females^7,8^. Individuals were further classified based on age estimation, stature, bone pathologies, and information derived from historical records. Long bones, dental and molar specimens were subsequently retrieved from the individuals included in this study, prioritizing those with optimal preservation.

After exhaustive historical research, the combination of anthropological studies, paleochemical analysis as radiocarbon or isotopic determination, the 3D scanning of the samples and posterior laser based mineral composition determination ending up with the genetic studies performed using Massive Parallel Sequencing (MPS), have been used to perform a comprehensive study of the individuals founded in the aforementioned crypt. Preliminary results identifying some of the individuals founded in the crypt have been already published ^7,8^.

This research seeks to resolve Columbus origin uncertainty through a comprehensive study of his documented direct descendants. By extracting and analysing autosomal DNA from these historical remains, we addressed two primary objectives: biogeographical ancestry estimation through ancestry informative markers (AIMs) and kinship analysis. The results presented herein are part of a broader, highly complex investigation into Columbus’s ancestry, providing a detailed genetic footprint of his lineage through his descendants’ autosomal genetic profiles.

This entire investigation has been conducted in accordance with all legal criteria and requirements regarding the preservation of the chain of custody and the legal validity of documentary and expert evidence ^9^.

## MATERIAL AND METHODS

### Historiography and documentation

A large amount of historical and genealogical data has been collected demonstrating that the skeletal remains analysed belong to the counts of Gelves lineage, direct descendants of Christopher Columbus. ^10-24^

### Sample’s Origin and Archaeological Context

An *in situ* anthropological study was performed on March 21st and 22nd 2022 at the crypt of the Church of Santa María de Gracia in Gelves, Seville. In the presence of a notary public, three wooden urns containing the remains of the Count of Gelves’ family were opened and documented via photography, 3D scanning, and osteometric measurements. To ensure transparency, impartial witnesses from the press were present during the exhumation^25^. The immediate collection of samples for DNA analysis made it possible to perform forensic genetics studies, with legal validity of an expert report.

### Physical Anthropology and Pathology ^26-42^

The minimum number of individuals (MNI) was determined using the White method. Biological sex was estimated following Bruzek (2002) and Buikstra & Ubelaker (1994). Age at death was assessed through multiple criteria: pubic symphysis morphology (Brooks & Suchey, 1990), auricular surface changes (Buckberry & Chamberlain, 2002), and sacral development (Passalacqua, 2009), supplemented by cranial suture closure (Meindl & Lovejoy, 1985) and dental wear (Brothwell, 1989). Stature and Body Mass Index (BMI) were estimated using standard osteometric formulae. Paleopathological analysis followed the clinical protocols of Ortner (2003) and Waldron (2008).

### Radiocarbon, Chemical and Isotopic Analysis ^43-58^

Prior to destructive sampling, 3D scans were generated using the Artec Space Spider and Artec Studio software to preserve the morphological integrity of the remains. Radiocarbon dating (C14) and stable isotope analysis (δ13C y δ15N) were conducted by Beta Analytic to determine chronologies and dietary patterns. Isobar Science performed strontium isotope analysis (^87^Sr / ^86^Sr) on dental enamel to assess geographical origin. Elemental composition was characterized via Laser-Induced Breakdown Spectroscopy (LIBS) using a SciAps Z-903 portable device, providing a "chemical fingerprint" of the remains.

### DNA extraction and quantification^59^

After cleaning and disinfection, bone samples were grinded using a Tissuelyser II (QIAGEN). Bone powder was incubated O/N in a lysis/decalcification buffer following previously described methods ^59^. Subsequent DNA extraction was carried out using the large volume protocol in an EZ2 Connect instrument (QIAGEN). DNA concentration process was carried out using Amicon®Ultra-0.5 columns (Millipore). The obtained DNA was quantified by real-time PCR using Quantifiler™Trio DNA Quantification kit in a QuantStudio5 thermocycler (ThermoFischer).

### Library preparation and targeted sequencing ^60^

Library preparation from genomic DNA was carried out following the ForenSeq® Kintellingence HT kit (QIAGEN) manufacturer’s protocol. DNAse/RNAse free water was used as negative amplification control and 1ng of the genomic DNA control sample NA24385 (HG002-Coriell Institute for Medical Research, Camden NJ) was included as positive control. Both were processed together with the test samples. To assess the size, quantity, and integrity, of the obtained libraries, these were quantified using both a fluorometric (Qubit™) and an electrophoretic method (TapeStation™). Subsequently, libraries were pooled together. The denaturalized pool was loaded on a MiSeq® FGx Reagent Cartridge. MPS run was set up in a MiSeq®FGx instrument. Targeted sequencing took place on a Standard MiSeq^®^FGx Flow Cell (QIAGEN).

### Data analysis

Sequenced libraries were visualised and analysed using the ForenSeq® Kintelligence Analysis Module in the ForenSeq® Universal Analysis Software (UAS) v2.7(QIAGEN). Allele calling was performed as described in the ForenSeq® Kintelligence Kit Reference Guide and the Universal Analysis Software – Kintelligence HT Module Reference Guide ^61,62^. All analysis within the UAS were performed with the default analytical and interpretation thresholds at 1.5 % in the ForenSeq® Kintelligence HT analysis method. This represents a minimum read count of 10 reads ^63,64^. Kinship calculations were carried out with the UAS associated local database, applying the manufacturer’s default settings.

### Pedigree Modelling and Biogeographical Ancestry Inference

Simultaneously, biogeographical ancestry (BGA) was inferred through a triangulated bioinformatic approach to ensure cross-validated results. Genomic datasets were analysed using Snipper v3.5 ^65^, and Genogeographer version 0.3.1 ^66,67,68,69,70^. Genealogical datasets were processed and modelled using the QuickPed analytical framework ^71^. Kinship coefficients across the expanded pedigree were systematically quantified using Henderson’s recursive algorithm, implemented through the kinship2 R package and the pedtools suite within the QuickPed framework ^72,73,74^. To evaluate the structural influence of the proposed genealogical configurations, a "Virtual Knock-out" analysis was performed. This involved a comparative assessment of kinship matrices under two mutually exclusive scenarios. Finally, ancestral nodes were scrutinized using Wright’s path coefficients to identify kinship intersections among descendants of a shared common ancestor ^75^.

## RESULTS

### Historiography and documentation

An analysis of historical documentation provided evidence that the familial pantheon is, in fact, the family tomb of the Counts of Gelves, which is indeed the place in the world where the largest number of direct descendants of Columbus are buried. Furthermore, it allowed to identify the historical figures buried there and provided their dates of birth and death, along with other key information needed to identify the individuals interred there.

### Archaeological and anthropological studies

Anthropological assessment determined the MNI at 12. The estimated ages at death and biological sex determined are shown in Table 1. The recovered data showed high concordance with the expected age and sex distribution, based on the historical records. Skeletal traits and pathologies were detected and registered in some of the individuals. The presence of metopism, a skeletal trait characterized by the persistence of the metopic suture in the skull, in three of the recovered individuals was remarkable.

**Table 1:**
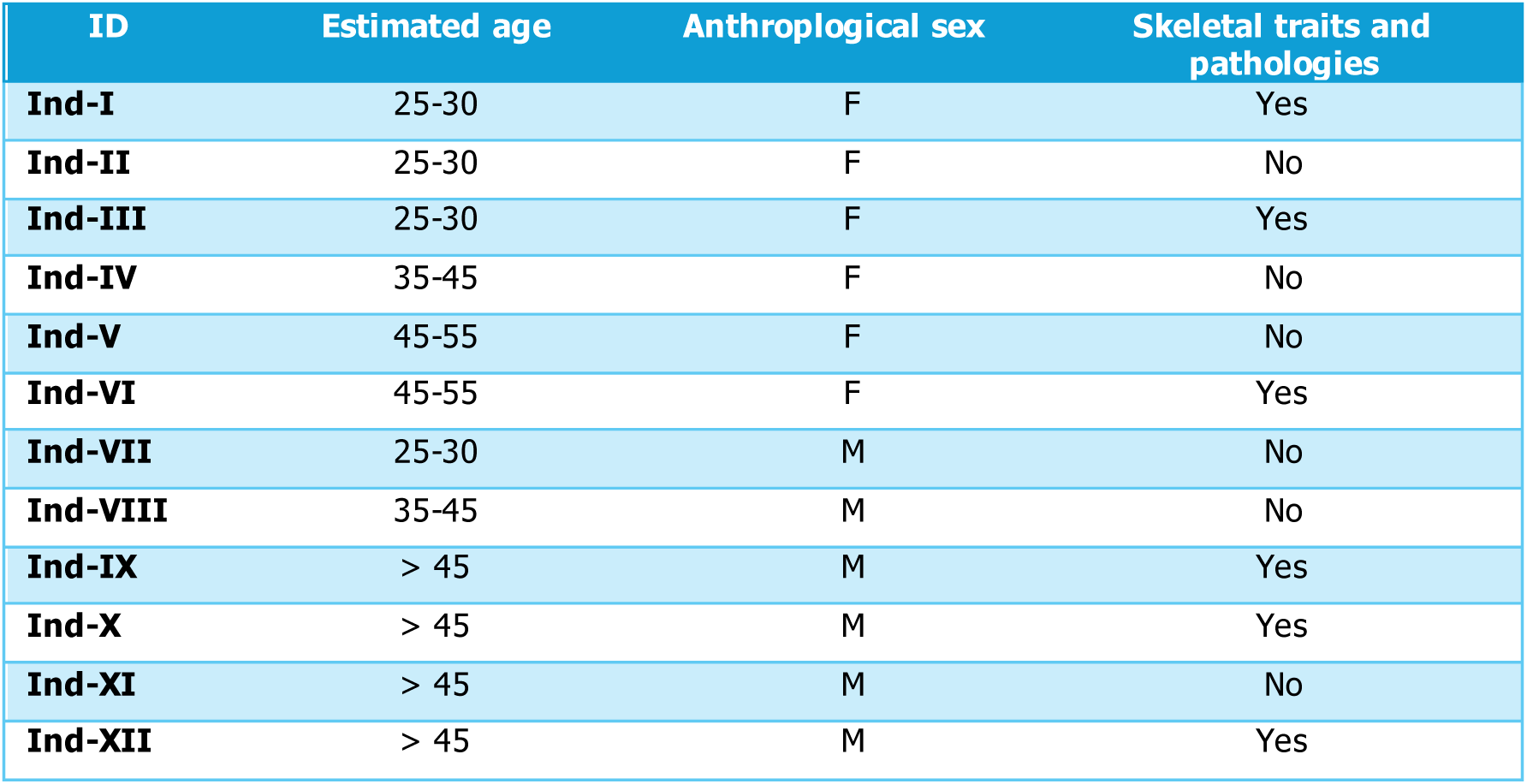
Individuals characterization through anthropological study. Estimated age at death; F: female; M: male.

| ID | Estimated age | Anthropological sex | Skeletal traits and pathologies |
| --- | --- | --- | --- |
| **Ind-I** | 25-30 | F | Yes |
| **Ind-II** | 25-30 | F | No |
| **Ind-III** | 25-30 | F | Yes |
| **Ind-IV** | 35-45 | F | No |
| **Ind-V** | 45-55 | F | No |
| **Ind-VI** | 45-55 | F | Yes |
| **Ind-VII** | 25-30 | M | No |
| **Ind-VIII** | 35-45 | M | No |
| **Ind-IX** | > 45 | M | Yes |
| **Ind-X** | > 45 | M | Yes |
| **Ind-XI** | > 45 | M | No |
| **Ind-XII** | > 45 | M | Yes |

LIBS detected traces of Al, Ti, Fe, Zn, Ba, and Pb, creating an elemental profile of each analysed skeletal sample, was used as a chemical fingerprint and allowed to ensure bone provenance and guaranteed the subsequent skeletal remains individualisation (data not shown). This characterization, together with the notary act, the photographs and 3D scan, provides absolute assurance of the chain of custody of the samples.

Samples from two individuals were subjected to studies of dietary and isotopic characterization. Stable isotope values obtained for the following individuals were as follows: a) Ind-I: IRMS δ13C: -17.7 ‰, IRMS δ15N: 12.6 ‰; b) Ind-II: IRMS δ13C: - 16.67 ± 0.30 ‰, IRMS δ15N: 13.05 ± 0.40 ‰. These values suggest a diet predominantly composed of marine proteins, which is consistent with the riverine and coastal environments of Gelves and Cádiz, where these individuals were residing.

### Genetic studies

To date, seven of the twelve individuals exhumated in this study have been genetically studied following amplicon library preparation protocols, with a large-scale SNP panel encompassing autosomal and Y- and X-chromosome markers. Up front, the genetically determined biological sex matched that previously determined on an anthropological basis.

The targeted SNP panel encompasses 56 Ancestry Informative Markers (AIMS) that have been analysed in order to establish the most probable biogeographical ancestry from the studied individuals. As expected, among the seven candidates, after applying several bioinformatic tools, all resulted compatible with European biogeographical ancestry (Figure 1).

**Figure 1.**
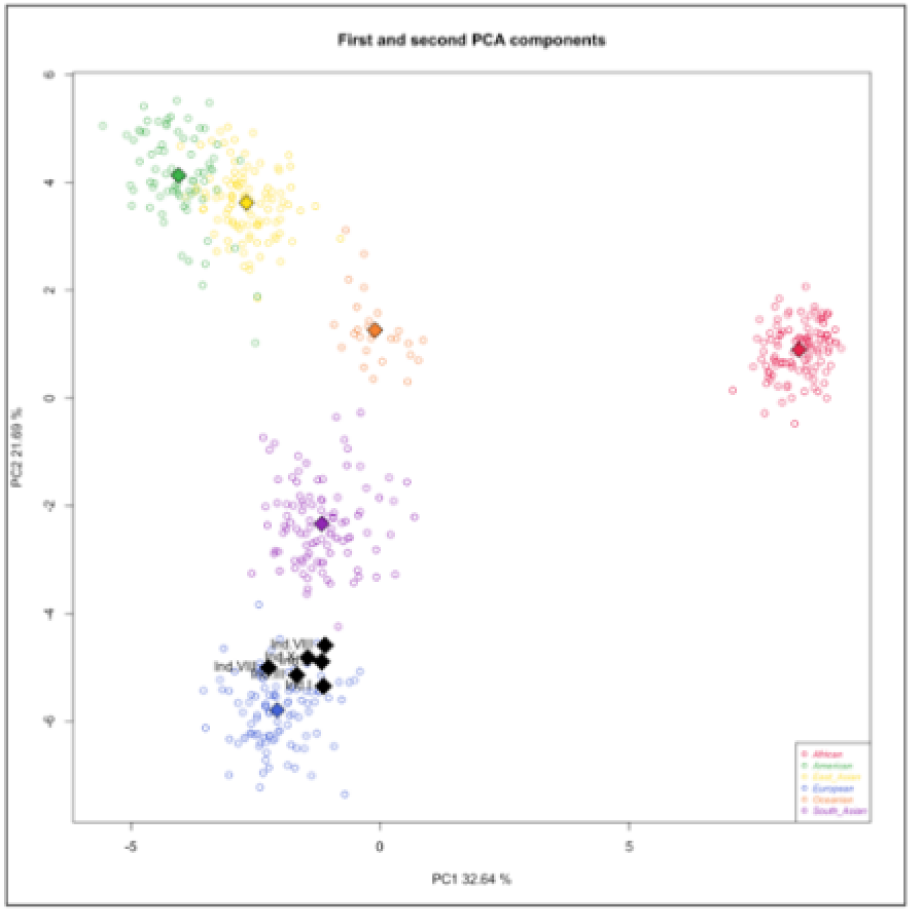
Principal Component Analysis (PCA) of genomic ancestry in the studied individuals. The scatterplot illustrates the projection of the first two principal components (PC1 and PC2) derived from th 56 autosomal SNPs. PC1 (32.6%) and PC_2 (21.7%) collectively account for 54.3% of the total genetic variation, capturing the primary axes of European population structure. The Gelves subjects (indicated by black diamonds]) exhibit a highly cohesive and compact clustering within the reference population. The tight spatial distribution of the samples in the PC_1/PC_2 plane underscores a robust genetic affinity and minimal individual outliers, providing strong empirical support for their biogeographical origin. Additional projections, including the third principal component PC3 = 7.94% (data not shown), demonstrate consistent clustering

Based on the documentary research, the only young adult male expected to be found in the crypt was Jorge Alberto de Portugal, the III Count of Gelves, who died at the age of 23. Ind. VII was the only male found whose age at death was compatible with his, thus it was classified as the most likely candidate for him. The specific data related to the comprehensive study done in this individual, including genetic profiling through autosomal, Y- and X-chromosome STR markers and SNPs, that allowed to confirm his identity have been already published ^8^

Subsequently, a genomic study based on Forenseq™ Kintelligence technology was carried out, which made it possible to establish kinship relationships between four of the individuals found in the crypt ^7^. These were individuals III, VIII, IX and X (Table 1). The documentary study carried out on the individuals present in the crypt together with the anthropological data and combined with the kinship results obtained in the aforementioned work, enabled the identification of these individuals and, consequently, their attribution to the corresponding historical figures (Table 2). Furthermore, the integration of all disciplines showed that individuals III, IX and X, who share a grandmother-son-grandson relationship, all exhibited metopism, an inherited condition, supporting even more the genetic based results.

**Table 2:** Identification of the historical characters: * Isabel de la Cueva was the first wife of the VI Count of Gelves (Pedro Nuño Colón de Portugal, a direct descendant of Columbus), and María de Castro Girón de Portugal was his second wife, whom he married after his first wife’s death. AA: anthropological estimated age; HA: Age at dead as determined by historical records; AS: Anthroplogical estimated biological sex; BS: Biological sex determined by genetic testing; SkT: Skeletal traits; C14: Radiocarbon based dating; LIBS: Laser-Induced Breakdown Spectroscopy; I: Isotopic analysis; DNA: Genetic study. Overall: Y stays for yes, meaning done; N stays for no, study not performed and ND for not determined yet.

| Ind. ID | AA | HA | AS | BS | SkT | C14 | IBS | I | DNA | Individual Identification | Lifespan |
| --- | --- | --- | --- | --- | --- | --- | --- | --- | --- | --- | --- |
| Ind-I | 25-30 | 32 | F | F | Y | [1539- 1635] | Y | Y | Y | ND | ND |
| Ind-II | 25-30 | 30 | F | F | N | [1628-1680] | Y | Y | Y | María de Castro Girón de Portugal, VI consort countess of Gelves * | 1640-1670 |
| Ind-III | 25-30 | 26 | F | F | Y | N | Y | N | Y | Isabel de la Cueva, VI consort countess of Gelves * | 1631-1657 |
| Ind-IV | 35-45 |  | F |  | N | N | Y | N | N | ND | ND |
| Ind-V | 45-55 |  | F |  | N | N | Y | N | N | ND | ND |
| Ind-VI | 45-55 |  | F |  | Y | N | Y | N | N | ND | ND |
| Ind-VII | 25-30 | 23 | M | M | N | Y | Y | Y | Y | Jorge Alberto de Portugal III count Gelves | 1566-1589 |
| Ind-VIII | 35-45 | 38 | M | M | N | N | Y | N | Y | Álvaro Jacinto Colón de Portugal, V consort count of Gelves | 1598-1636 |
| Ind-IX | > 45 | 55 | M | M | Y | N | Y | N | Y | Pedro Manuel Florentín Colón de Portugal y Ayala, VIII count of Gelves | 1676-1736 |
| Ind-X | > 45 | 56 | M | M | Y | N | Y | N | Y | Pedro Manuel Colón de Portugal y de la Cueva, VII count of Gelves | 1651-1710 |
| Ind-XI | > 45 |  | M |  | N | N | Y | N | N | ND | ND |
| Ind-XII | > 45 |  | M |  | Y | N | Y | N | N | ND | ND |

Finally, another kinship came to light after further genetic studies. This relationship was established between Individual VII, previously identified as Jorge Alberto de Portugal, and Individual II, a woman who was determined as 25–30 years old at the time of her death. According to historical documents and anthropological findings, there were two female candidates who matched to this age: Individual II and Individual I. In order to rule out which sample could correspond to each woman, samples from individuals I and II were sent for C14 analysis, what made it possible to date both women and thus distinguish them in time.

For Ind.I, the date of death was determined with the highest probability falling within the years 1539 to 1635. On the other hand, in the case of Ind.II, the most likely estimated range for her date of death was determined between 1628 and 1680, showing that individual Ind.I dates from approximately one century earlier than Ind.II.

Once more, putting together the data from all disciplines involved in this work, the identification of individual II was possible, according to the anthropological estimated age of dead, the radiocarbon datation, the chemical footprint and the historical records, showing that Individual II corresponds to María de Castro Girón de Portugal, VI consort countess of Gelves.

A summary of the studies and findings relating to the individuals in the crypt is shown in Table 2.

### Kinship analysis

The UAS v 2.7 allows to do kinship calculations based on the SNPs tested with the Kintelligence HT panel. The results obtained for Ind.VII and Ind.II are shown in Table 3

**Table 3:** Kintelligence HT results. . Total reads: number of reads per sample; Total SNPs: number of SNPs typed; X/Y-SNPs: SNPs located in X/Y Chromosome; aSNPs: ancestry SNPs; phSNPS: phenotype SNPs; iSNPs: identity SNPS and kSNPs: kinship SNPs. Typed markers are expressed as total number (Nr) and percentage of markers present in the panel (%) typed.

|  | Total reads |  | Total SNPs |  | X-SNPs |  | Y-SNPs |  | aSNPs |  | phSNPs |  | iSNPs |  | kSNPs |  |
| --- | --- | --- | --- | --- | --- | --- | --- | --- | --- | --- | --- | --- | --- | --- | --- | --- |
| Sample ID | Nr |  | Nr | % | Nr | % | Nr | % | Nr | % | Nr | % | Nr | % | Nr | % |
| Ind.II | 1.922.632 |  | 9.385 | 92,51 | 104 | 98,11 | 0 | 0 | 48 | 85,71 | 24 | 100 | 84 | 89,36 | 9127 | 92,50 |
| Ind. VII | 2.157.352 |  | 9.469 | 92,56 | 98 | 92,45 | 76 | 89,41 | 48 | 85,71 | 24 | 100 | 90 | 95,74 | 9135 | 92,58 |

**Table 4:** Kinship study results. SNP overlap: number of SNPs in common between two samples; Shared DNA: Total amount of shared DNA between two samples. cM: Centimorgans; Longest shared segment: The longest stretch of shared DNA between two samples.

| Sample ID | Related Sample ID | SNP overlap | Shared DNA (cM) | Longest shared segment (cM) | Kinship coefficient |
| --- | --- | --- | --- | --- | --- |
| Ind.II | Ind. VII | 9109 | 123,966 | 50,605 | 0,014 |

The main results obtained out of the UAS local database algorithm related to the detected kinship between the two individuals identified as Jorge Alberto de Portugal, 3^rd^ count of Gelves (Ind.VII) and María de Castro Girón de Portugal, VI countess consort of Gelves (Ind. II) are shown in the following table:

The initially unexpected biological kinship between María de Castro and Jorge Alberto de Portugal necessitated a rigorous multi-generational analysis. While her interment in the crypt was historically attributed to her status as the consort of the VI Count of Gelves rather than direct consanguinity, the genetic evidence prompted an exhaustive investigation into her full ancestry. This research stated on her high-status paternal lineage as the daughter of the IX Count of Lemos (noble house of Galicia, Spain) but encompassed a full reconstruction of both maternal and paternal branches to ensure a comprehensive screening of the pedigree.

Systematic mapping of the shared ancestral landscape identified a profound convergence within two prominent noble houses: Zúñiga (Navarra) and Sotomayor (Galicia). These lineages, both originating from Northern Iberia, were found to exert a consistent influence on the shared genomic inheritance between the subjects. Pedigree reconstruction identified Pedro Álvarez de Sotomayor—a noble descendant of both houses—as a high-centrality node. His recurrent presence across multiple ancestral pathways within the expanded tree provides the requisite connectivity to explain the identified kinship, effectively functioning as a very likely biological nexus for the transmission of the ancestral genetic signal.

Consequently, based on the familial pedigrees, two competing hypotheses were formulated for in silico statistical evaluation: (a) the observed kinship is derived from Pedro Álvarez de Sotomayor as the second-great-grandfather of Jorge Alberto de Portugal; versus (b) the null hypothesis, in which the second-great-grandfather of Jorge Alberto de Portugal is an individual possessing no biological affinity with Pedro Álvarez de Sotomayor, and thus has no influence on the biological relationship detected in the genetic study (Figure 3).

**Figure 2:**
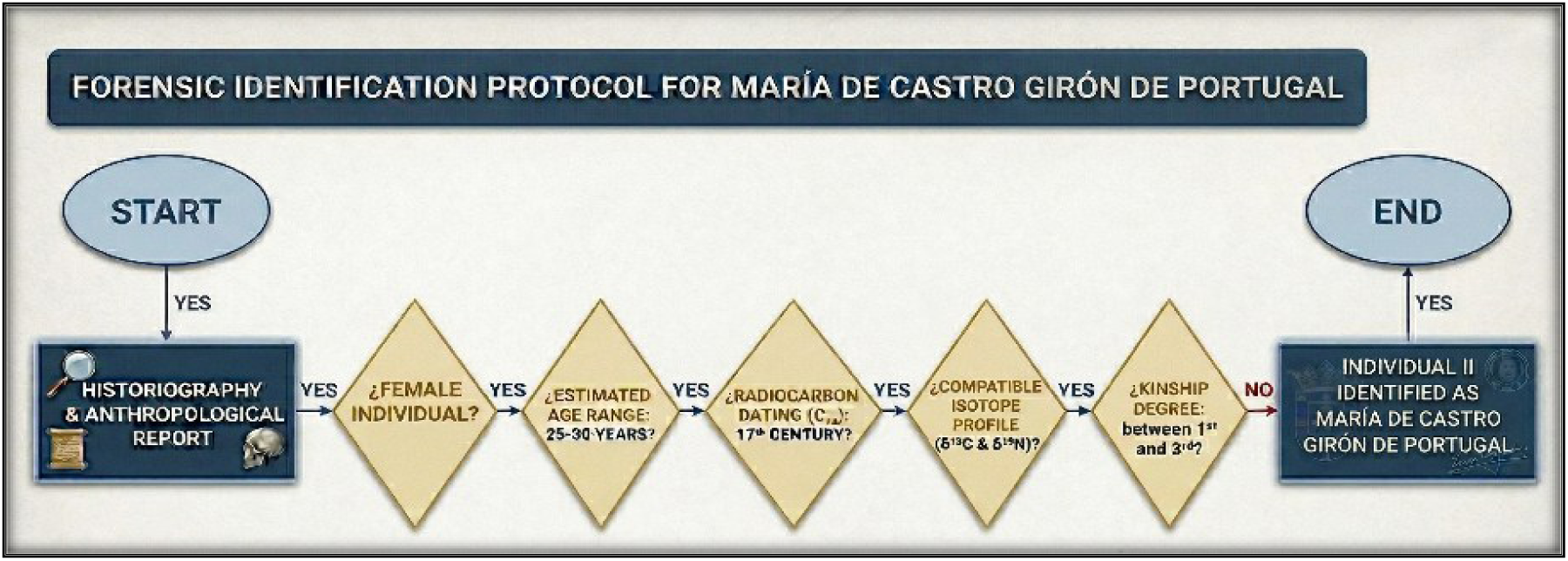
Forensic identification protocol for María de Castro Girón de Portugal.

**Figure 3.**
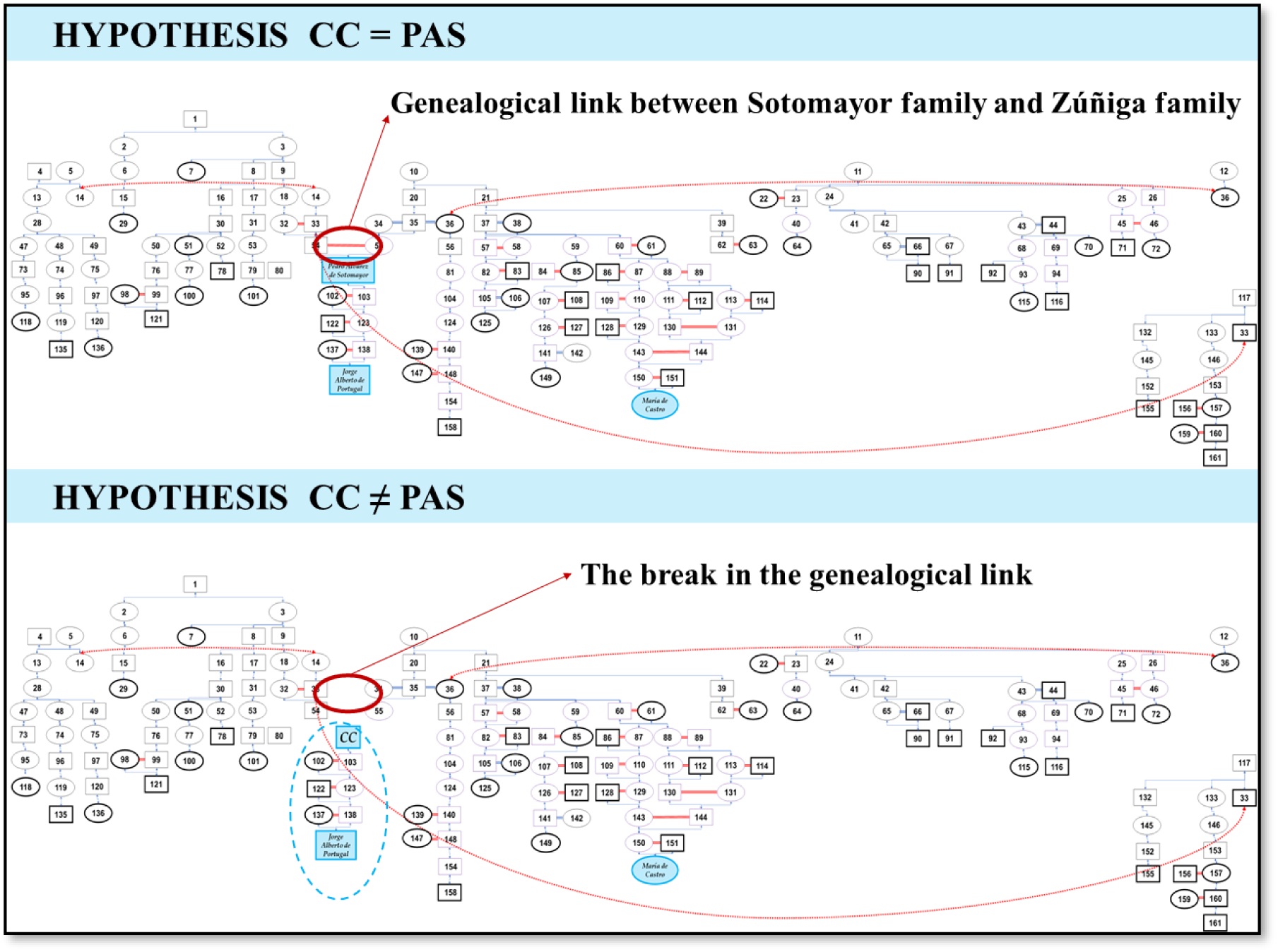
Pedigrees representing the two postulated hypotheses. Individuals are represented by numbers. Squares represent male individuals; circles represent female individuals. Marriages shown in red indicate consanguinity. Red doted arrows show relationships from the same individuals with other family branches. CC: Christophorus Columbus; PAS: Pedro Álvarez de Sotomayor

The resulting variance between these matrices delineated the estimated net genetic contribution of Pedro Álvarez de Sotomayor to the kinship detected between María de Castro and Jorge Alberto de Portugal, in strict accordance with Mendelian inheritance patterns. The identification of multiple nodes with equivalent contribution values served to validate a sustained "genealogical tunnel"— a constant connecting lineage demonstrating that the relatedness between distant cousins is fundamentally contingent upon this specific chain of individuals.

Both hypotheses were analysed using the R package pedtools (by Vigeland). This analysis characterized the genetic relationship between individuals Jorge Alberto de Portugal and María de Castro within a deep 16-generation pedigree, encompassing the Zuñiga and Sotomayor ancestors. Despite a significant temporal distance, a detectable kinship remains (Φ = 0.00243). These findings identified Pedro Álvarez de Sotomayor as a pivotal genetic bridge; the exclusion of this node, representing hypothesis b, resulted in a total loss of kinship (r = 0), defining Pedro Álvarez de Sotomayor as an obligatory ancestor for the observed IBD (Identical By Descent) sharing.

## DISCUSSION

The integration of high-resolution genomic data with forensic anthropology and multi-generational pedigree modelling presented in this work, has provided a transformative perspective on the ancestral origins of Christopher Columbus. By analysing the genetic legacy bequeathed to his direct descendants, this study successfully projects the lineage backward to identify his biological roots.

The empirical identification of a 50.605 cM contiguous IBD segment between María de Castro Girón de Portugal (Ind. II) and Jorge Alberto de Portugal (Ind. VII)—the III Count of Gelves and great-great-grandson of Columbus—, prompted a novel line of inquiry. This shared genetic signal, identified within an archaeological context where direct relationship was not historically documented, served as the primary catalyst for the multi-layered genomic and genealogical investigation presented here. By integrating high-resolution autosomal data with forensic pedigree modelling, this study establishes a definitive framework for re-evaluating the ancestral origins of Christopher Columbus. Therefore, a comprehensive pedigree based on primary sources ranging from the 11th to the 18th century was performed. This multi-secular investigation aimed to map the specific points of convergence between the lineages and to identify the origin of the unexpected biological nexus.

The crucial discovery of this research is the role of Pedro Álvarez de Sotomayor within the consolidated pedigree. Systematic modelling corroborated the empirical results obtained by identifying a significant kinship surplus between María de Castro and Jorge Alberto de Portugal, that exceeded documented genealogical predictions. Based on the genealogical research, the present study demonstrates that, to our knowledge, Pedro Álvarez de Sotomayor is the unique ancestor capable of reconciling this genomic covariance.

Moreover, computational exclusion via the "Virtual Knock-out" technique confirms Pedro Álvarez de Sotomayor as the critical node in this biological bridge. Upon his removal from the expanded tree, the kinship between María de Castro and the second-great-grandson of Columbus is eliminated. Across 16 generations of scrutinized connections, no other individual was identified who could provide the specific genetic inheritance required to explain the identified kinship.

These findings definitively point to the ancestral origin of the Columbus lineage within the noble houses of Northern Spain, specifically the Zúñiga (Navarre) and Sotomayor (Galicia) families.

## CONCLUSION

By synthesizing MPS-derived autosomal profiles with rigorous historiographical and anthropological data, this study establishes a coherent framework linking the Columbus lineage to the elder Galician and Navarrese nobility. The results confirm that the remains deposited in the church of Santa María de Graciás crypt (familial pantheon of the counts of Gelves) are indeed those of the counts of Gelves, as historical accounts had suggested. Furthermore, they are direct descendants of Christopher Columbus and share a common genetic architecture with the Sotomayor and Zúñiga houses. Collectively, these findings provide for the first time robust genetic support for the hypothesis of a Galician provenance for Christopher Columbus, laying a definitive foundation within the scientific discourse for the re-evaluation of his historical identity.

## ACKNOWLEGEMENTS

The authors express their gratitude to the Archdiocese of Seville, specifically Bishop Teodoro León Muñoz, for the authorization that made this research possible. We thank Alba Hernández for her accurate laboratory work, and Ana María Ortiz Paredes of "El Mundo" for her professional chronicling of the archaeological process.

## AUTHORSHIP CONTRIBUTION

Sample collection and "in situ" archaeological work: Yravedra Saez de los Terreros, J, Bonilla A, Tirapu M, Albert M, Jímenez P, Herránz D. Legal assistance, organization and independent sponsorship of the scientific project: García, M.C. Anthropologic study: Tirapu M. DNA extraction, genetic studies and quality control during analyses: Navarro-Vera, I.

